## supplemental information for "Differential effects of redox conditions on the decomposition of litter and soil organic matter"

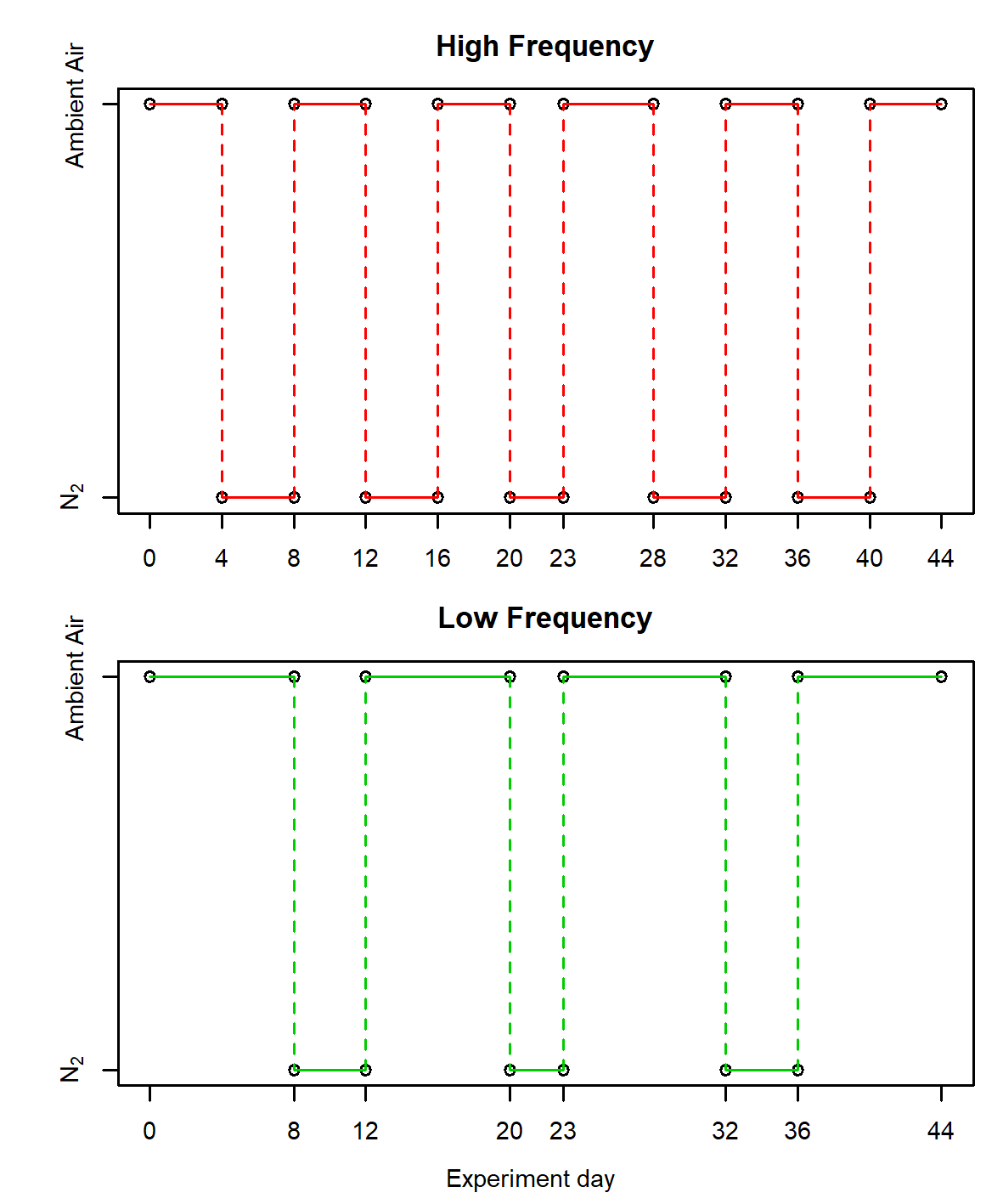

Fig. S1. Changes in headspace composition for samples from the two redox fluctuating treatments.

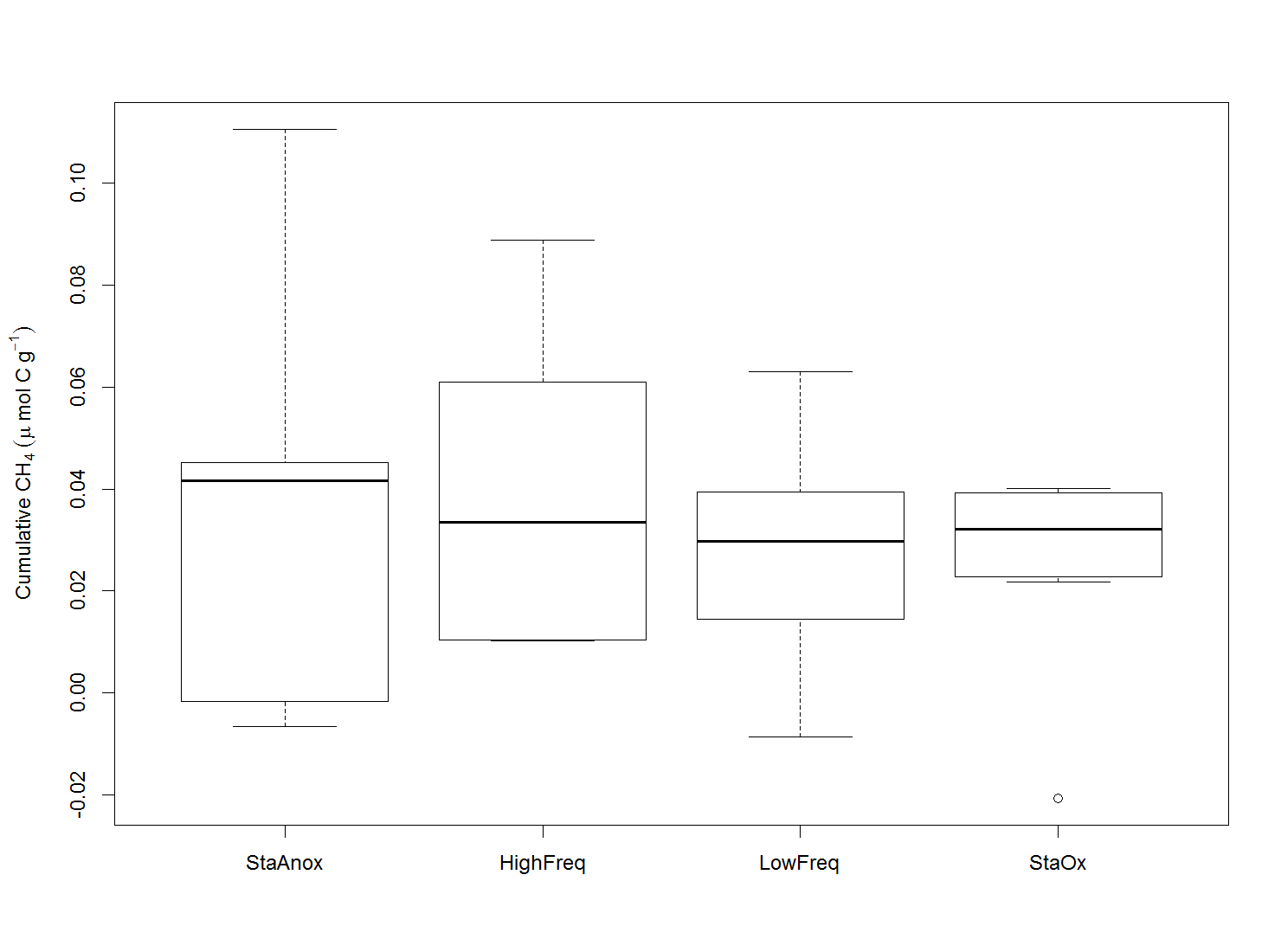

Fig. S2. Cumulative CH_4_ production over the incubation experiment.

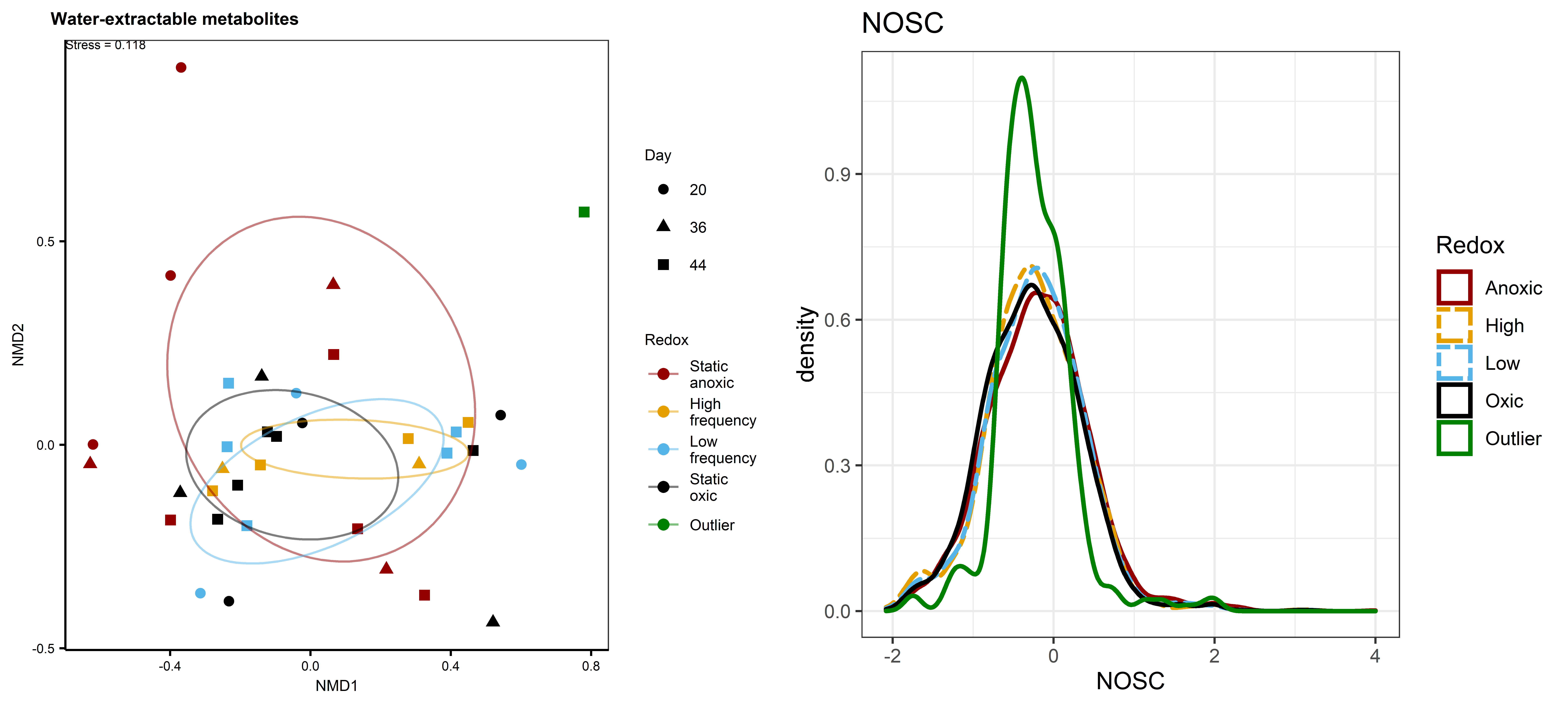

Fig. S3. (left) Non-metric multidimensional scaling (NMDS) plot comparing the composition of water-extractable organic matter between the outlier and other samples. Data were derived from FTICR-MS analysis. The eclipse indicates the standard deviation of each redox treatment. Note the deviation of outlier from rest of the samples. (right) The probability density curves of NOSC values of water-extractable organic matter comparing the outlier with other samples. Note the large difference between the outlier and other samples and the small differences among redox treatments.

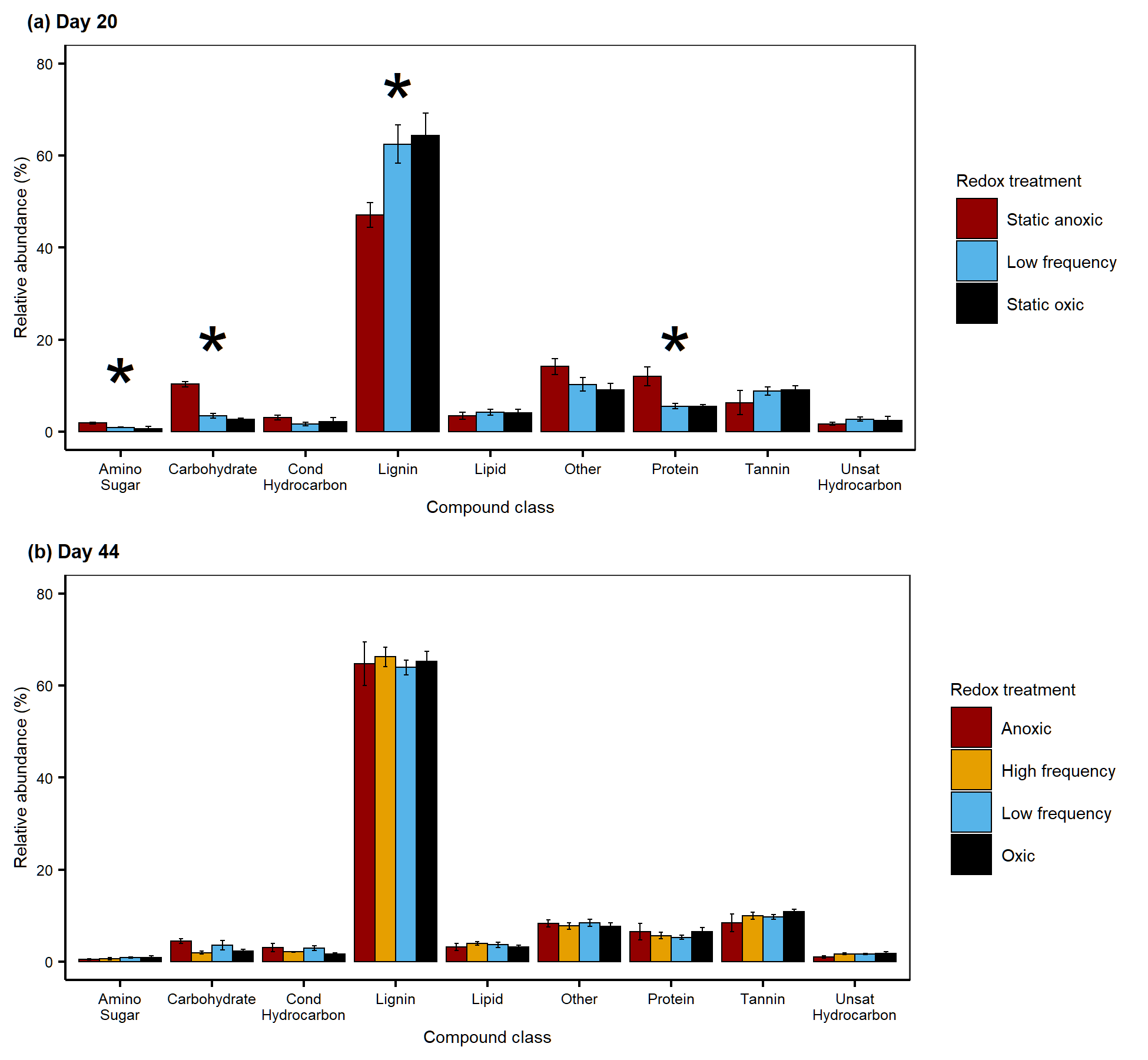

Fig. S4. Effects of four redox treatments on the relative abundances of chemical compound classes extracted by water on day 20 (a) and day 44 (b) of a tropical soil incubation. Data were derived from FTICR-MS analysis. Error bars indicate standard errors of means. * indicate significant effects of redox treatments (ANOVA) at α = 0.05 level. On days 20 and 44, *n* = 3 and 5 per treatment, respectively. High frequency treatment from day 20 was not included. Two outliers were removed from day 44. See the Materials and Methods for details.

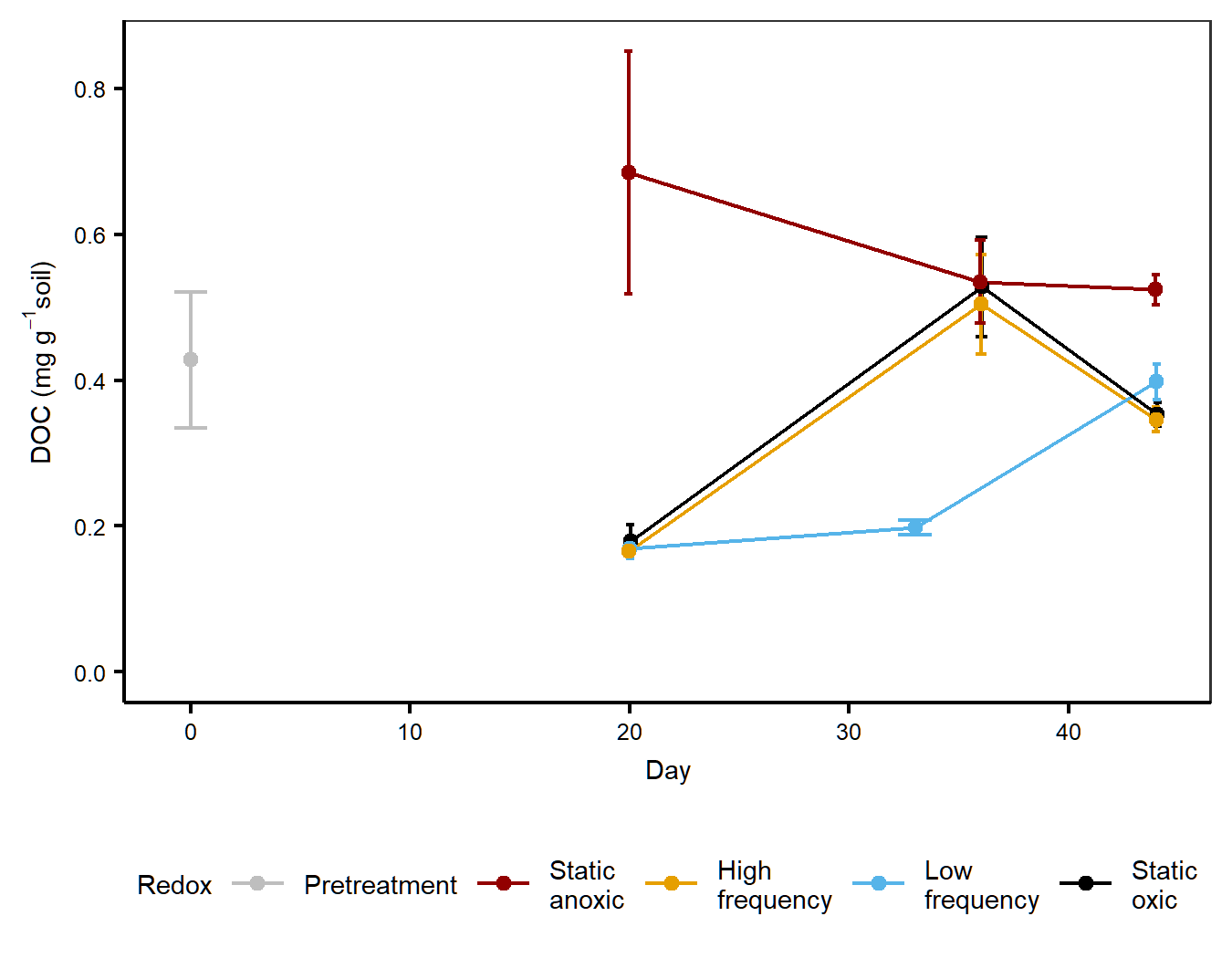

Fig. S5. Effects of redox treatments on the concentrations of water-extractable dissolved organic C (DOC). Pretreatment values are in grey. *n* = 3 per timepoint and treatment except *n* = 5 on day 44.

Table S1. Carbon concentration and ^13^C abundance of background soil and ryegrass litter (Mean ± S.E.).

|  | C concentration | ^13^C abundance |
| --- | --- | --- |
| Background soil (n = 3) | 6.1 ± 0.4% | -29.0 ± 0.8 ‰ |
| Ryegrass litter (n = 6) | 41.0 ± 0.1% | 96.7 ± 0.1 atom% |

Table S2. Total number of water-extractable molecular formulas identified by FTICR-MS analysis per sampling days and treatments.

| Sampling Day | Static anoxic | High frequency | Low frequency | Static oxic |
| --- | --- | --- | --- | --- |
| 20 | 3795 | 584^a^ | 2635 | 2499 |
| 36 | 3070 | 1786 | N/A^b^ | 2888 |
| 44 | 3938 | 3394 | 4364 | 4697 |

^a^ Treatment was removed from further analysis due to the extremely low number of peaks identified.

^b^ Treatment was not included in the original experimental design.
